# AstraKit: Customizable, reproducible workflows for biomedical research and precision medicine

**DOI:** 10.64898/2026.01.25.701622

**Authors:** Nadine S. Kurz, Kevin Kornrumpf, Martin Stoves, Jürgen Dönitz

## Abstract

**Motivation:** The success of precision medicine and biomedical research depends on the availability of efficient software solutions for processing and interpreting genetic variants, interpreting multi-omics data, and integrating drug screen analyses. However, fragmented bioinformatics tools compel researchers and clinicians to resort to error-prone manual pipelines.

**Results:** We present AstraKit, a unified KNIME workflow suite enabling end-to-end precision medicine analytics. AstraKit introduces three transformative innovations: 1) Dynamic variant interpretation with customizable annotation and filtering for disease-specific genomic contexts; 2) Multi-layered omics analyses integrating genomic, transcriptomic, and epigenetic data; and 3) Translational drug matching that correlates in vitro drug screens with clinical outcomes. Validated across oncology cohorts, AstraKit demonstrates concordance between experimental drug sensitivity and clinical outcomes, resolving discordances to uncover resistance mechanisms. By unifying variant analysis, multi-omics, and drug response modeling on a single customizable platform, AstraKit eliminates siloed workflows, accelerating biomarker validation and enabling clinicians to directly link molecular profiles to therapeutic decisions. As all AstraKit workflows are open-source and platform-independent, we provide a versatile comprehensive software suite for a multitude of tasks in bioinformatics and precision medicine.

**Availability and implementation:** The KNIME workflows are available at KNIME Hub https://hub.knime.com/bioinf_goe/spaces/Public/AstraKit~lfVsGBY2HnPYc1h1/. The source code is available at https://gitlab.gwdg.de/MedBioinf/mtb/astrakit.

## 1 Introduction

Precision medicine demands robust computational frameworks to transform heterogeneous molecular data into clinically actionable insights. While nextgeneration sequencing (NGS) and high-throughput drug screening have exponentially expanded our capacity to generate genomic, transcriptomic, epigenomic, and functional drug response data, the lack of integrated, end-to-end analytical pipelines remains a critical barrier to reproducible and clinically translatable research. Software packages typically operate in silos: variant calling (GATK [1]), genomic operations (samtools [2], bedtools [3]), annotation (ANNOVAR [4], VEP [5], Onkopus [6]), and filtering (SNPSift [7], AdaGenes [8]) address isolated steps in the analytical continuum. Consequently, researchers must manually chain tools across incompatible data formats (VCF [9], MAF, BED), perform auxiliary tasks like genome assembly coordinate conversion (LiftOver), and implement custom filtering criteria—compromising traceability, reproducibility, and accessibility for non-programmers. This fragmentation impedes two critical translational needs: First, robust cohort-level variant analysis, essential for identifying recurrent mutations and clinical associations, often requires ad hoc integration of specialized tools (e.g., *maftools* [10]) with upstream processing, hindering scalable biomarker discovery. The joint analysis of various omics data, including transcriptomics and epigenetics, places further demands on data integration. Second, while clinical interpretation tools (e.g., MTB-Report [11], Pandrugs [12]) suggest targeted therapies from mutational profiles, and drug screen platforms (e.g., Breeze [13]) enable in vitro data exploration, no existing framework systematically links clinical genomic classifications to functional drug response validation.

Workflow systems like Snakemake [14] and Nextflow [15] improve reproducibility but require coding expertise, while cloud-based platforms like Galaxy [16] may pose challenges for sensitive clinical data due to privacy regulations. Although the Jupyter environment [17] enhances codes reproducibility, it lacks native workflow orchestration and standardized sharing mechanisms for complex pipelines. KNIME [18] provides a local, GUI-driven environment for workflow assembly. Prior workflow toolkits for precision medicine focused narrowly on variant calling/annotation [19, 20, 21, 22] or reference genome processing [23], without unifying variant interpretation, multiomics integration, or drug response validation. While bioinformatic workflow managers offer reproducibility, scalability, and portability [24, 25], the potential for dynamically combining heterogeneous omics data with clinical evidence and functional validation—critical for precision oncology—remains largely unrealized.

Here we present AstraKit, a comprehensive suite of modular, scalable, and reproducible workflows supporting a multitude of processing steps in bioinformatics and precision medicine. Unlike existing tools, AstraKit provides integrated workflows spanning from molecular profile processing and analysis to clinical interpretation and experimental validation. It supports seamless, format-agnostic data preprocessing, rule-based variant annotation/filtering, and cohort-level analysis. Critically, AstraKit bridges three previously disconnected domains: Multi-omics integration for joint analysis of mutations, gene expression, and epigenetic data—enabling pathway enrichment, survival modeling, direct linkage of clinical variant interpretations to drug sensitivity profiling, where patient-derived genomic biomarkers are systematically validated against high-throughput drug screen data (e.g., IC_50_, GR metrics) to identify actionable therapeutic targets. By unifying computational genomics with functional validation in a single framework, AstraKit accelerates the translation of molecular insights into clinically testable hypotheses.

## 2 Materials and Methods

### 2.1 Datasets and analytical workflows

To test and validate the workflows for data preprocessing and variant interpretation, we used publicly available datasets, including somatic variants in VCF format [26, 27]. For the variant data cohort analysis, we downloaded somatic variant data, gene expression, and DNA methylation files from 32 cancer types in The Cancer Genome Atlas (TCGA) using the TC-GAbiolinks package [28]. TP53 tumor variants were downloaded from the TP53 database [29]. Drug sensitivity data was retrieved from cancer cell line data from DepMap [30] and GDSC [31] (release 8.5), which includes IC_50_ and AUC values from drug sensitivity assays. The AstraKit framework was constructed using the KNIME analytics platform (v5.8.0) using Python (v3.11) and R (v4.5.2) extensions.

### 2.2 Variant annotation and interpretation

Data parsing and genome assembly conversion were implemented using *AdaGenes* [8], which supports VCF, MAF, FASTA, CSV, and plain text formats, as well as LiftOver between GRCh37/GRCh38/T2T-CHM13 assemblies. For variant annotation and interpretation, we developed custom KNIME nodes leveraging the *Onkopus* Python package [6]. Annotated variants were serialized as pickled objects for internode transfer. Genetic variants at transcript/protein levels were mapped to genomic coordinates using `SeqCAT` [32]. To standardize cancer type nomenclature across heterogeneous sources, we retrieved targeted therapy evidence from CIViC [33], MetaKB [34], and OncoKB [35]. Cancer entities were mapped to the OncoTree ontology [36] via a two-step process:1) Exact string matching against OncoTree’s canonical cancer type names, and 2) fuzzy matching for unmatched entries, selecting the highest-scoring OncoTree term. The token sort ratio normalizes Levenshtein distance by string length by first tokenizing,sorting, and rejoining words to handle word-order variations:

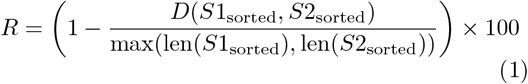

Where *S*1*sorted* and *S*2*sorted* are token-sorted versions of the input strings. Drug names were standardized to ChEMBL IDs [37] by resolving synonyms via the ChEMBL Onkopus API, ensuring consistent aggregation of drug response data.

### 2.3 Cohort and multi-omics analyses

Cohort-specific analyses and visualizations were implemented using maftools, scikit-learn [38], and seaborn [39]. Protein structural analyses leveraged the Onkopus Application Programming Interface (API) to derive 3D features. Biochemical properties were computed using *BioPython* [40], and AA*G* (change in folding free energy) was predicted with *FoldX* (v5.0) [41] via the `BuildModel` command:

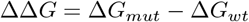

### 2.4 Drug screen analysis

To analyze drug sensitivity in relation to clinical outcomes, IC_50_ values were first converted to molar concentration (M) based on user-specified input units (e.g. nM, *μM)*. IC_50_ values were then transformed to pIC_50_ values:

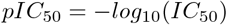

This logarithmic transformation linearizes the concentration-response relationship and stabilizes variance, with higher pIC_50_ values indicating greater drug sensitivity (i.e., lower molar IC_50_). For samplewise normalization (to account for inter-sample variability in assay performance), we implemented a robust percentile-based approach within each sample (grouped by sample ID). Normalization was performed directly on the pIC_50_ scale to preserve the linearized relationship:

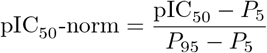

where *P*_5_ and *P*_95_ represent the 5th and 95th percentiles of pIC_50_ values within the sample-specific drug response distribution. This scaling maps the central 90% of values to [0, 1], with values below *P*_5_ assigned 0 (maximal resistance) and those above *P*_10_ and those above *P*_95_ assigned 1 (maximal sensitivity). For binary classification, normalized values were categorized using a user-defined threshold, defaulting to 0.5:

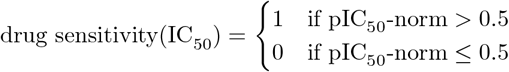

This threshold defines sensitivity as responses in the upper half of the normalized scale (i.e., above the midpoint between *P*_5_ and *P*_95_), not the raw median. Note that 0.5 corresponds to the scaled midpoint of the central 90% distribution, which may differ from the true median if the distribution is skewed. The predicted in vitro sensitivity was then compared against clinical response data. Clinical outcomes *r*_*i*_ were defined per standard criteria, where:

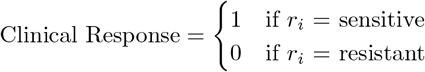

To quantify agreement between predicted and clinical responses, we evaluated binary classification performance using receiver operating characteristic (ROC-AUC).

## 3 Results

### 3.1 The AstraKit workflow suite

AstraKit delivers a comprehensive suite of modular, automated workflows for a multitude of tasks in precision medicine and biomedical research, including genomic data processing, multi-omics integration, and drug screen analysis (Figure 1). These pipelines support end-to-end genetic data analysis—from raw sequencing data through variant calling, molecular data preprocessing, annotation, filtering, and clinical interpretation. They further facilitate robust cohortlevel analyses and adaptive integration of heterogeneous omics data. Critically, AstraKit introduces innovative tools for preclinical model assessment and clinical trial evaluation by integrating molecular data with experimental drug screen results. Implemented in KNIME, AstraKit provides a graphical user interface that allows researchers to customize, modify, and execute workflows without coding expertise. Built-in workflow saving ensures reproducibility and seamless sharing of research results.

**Figure 1:**
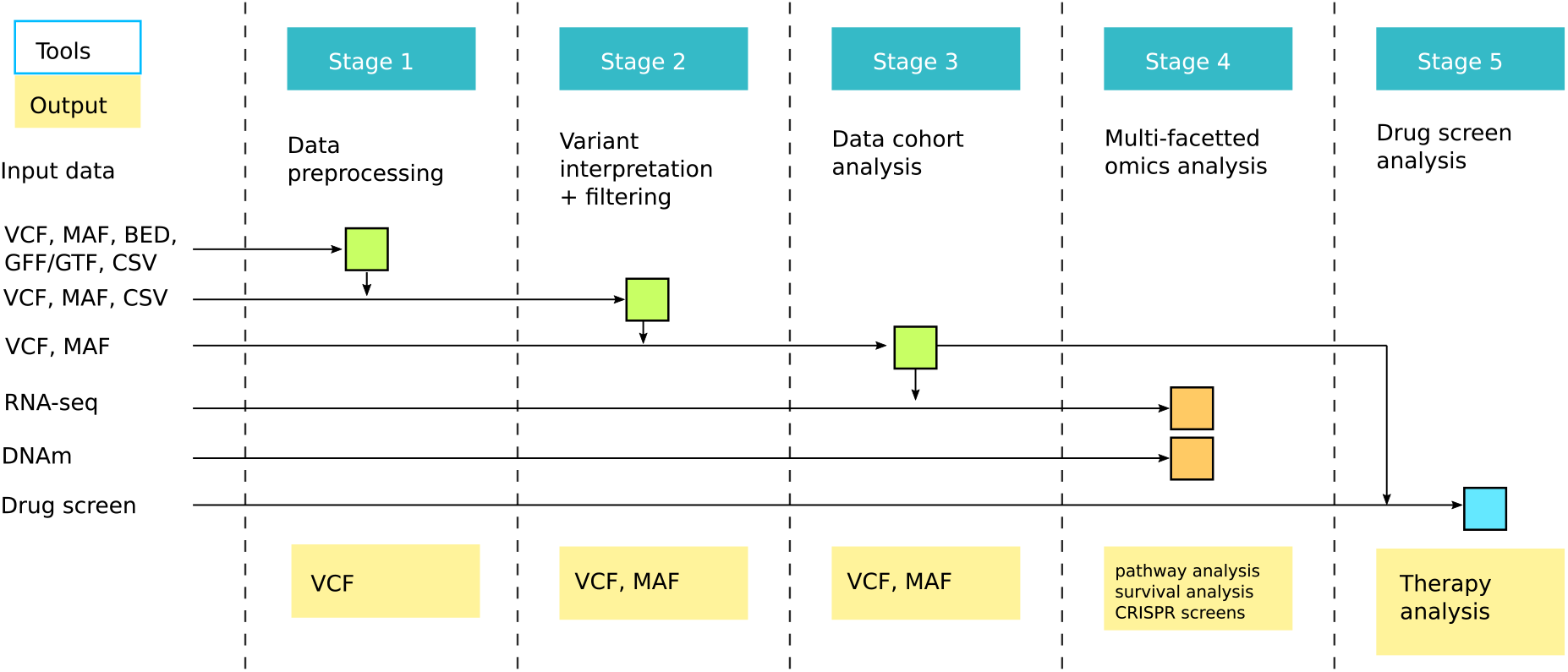
Overview of the AstraKit scope. AstraKit is able to preprocess mutational data in various formats, including VCF, MAD, BED, GFF/GTF, and CSV, for file format conversions and genome assembly coordinate conversion (LiftOver). Variant interpretation workflows support modular annotation and filtering using customizable rules. Cohort analysis leverages tools like *maftools* for mutation pattern detection, while multi-omics analyses facilitates joint analysis of genomic, transcriptomic, and epigenomic data. Additionally, drug sensitivity validation is performed through integrated drug screen analysis workflows.

### 3.2 Customizable mutational profile processing

AstraKit’s variant interpretation framework empowers users to construct annotation and filtering pipelines through a unique modular architecture. Each processing step (e.g., LiftOver, annotation, filtering) is encapsulated in standalone, interoperable nodes. This modularity is enabled by a standardized branched JSON data structure that serves as the universal input/output format across all nodes. By enforcing consistent data handling, nodes can be freely combined without manual data reformatting to generate reproducible, context-specific workflows. The suite provides extensive adaptable nodes for annotation (e.g., pathogenicity predictors, protein features, clinical evidence), filtering, LiftOver, and protein-to-genome conversion (resolving coordinate system mismatches between protein and genomic variants). Workflows dynamically begin with format-specific readers (VCF, MAF, CSV) and conclude with export nodes, supporting flexible pipeline design. For instance, a workflow can conduct a functional annotation of variant cohorts (Figure 2A), where protein-level variants are mapped to genomic coordinates, merged, and annotated with functional context—including transcript isoforms, protein surface accessibility, alpha-carbon distances, and secondary structures—to reveal mutation-driven mechanistic insights into disease pathogenesis. For clinical interpretation, a workflow (Figure 2B) may begin by annotating with population allele frequencies (gnomAD), then filtering common variants before layering pathogenicity predictions (e.g., SIFT, PolyPhen), protein structural data, and ClinVar classifications. Benign/non-coding variants are iteratively excluded, enabling efficient prioritization of clinically actionable alterations. Final outputs include annotated VCFs, column-wise variant reports (Figure 2C), and evidence-based therapy recommendations ranked by clinical actionability (Figure 2D), facilitating seamless transition from analysis to clinical decision support.

**Figure 2:**
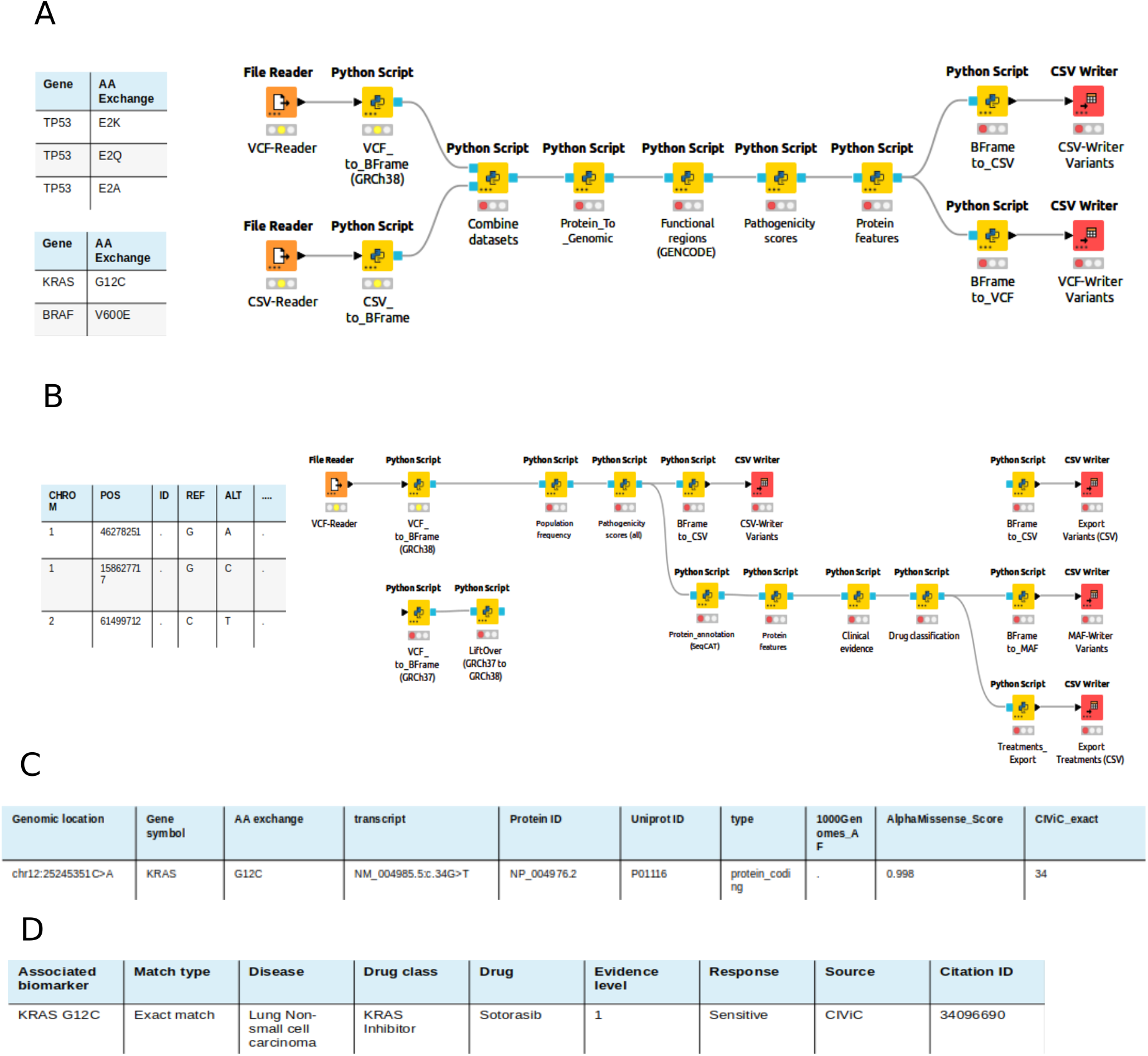
Customizable variant interpretation workflows in AstraKit. (**A**) A KNIME variant annotation workflow start with reading in one or multiple data files. Users can then dynamically align variant processing nodes to annotate the variants with selected features. The annotated variants can be written out again in different data formats. (**B**) For clinical interpretation, AstraKit provides the functionality to add clinical evidence and drug classification annotation, and the option to export therapeutic options for a patient. (**C**) AstraKit can export the functional annotations of variants, as well as (**D**) the retrieved evidence-based biomarkers for selected targeted therapies.

### 3.3 Cohort and multi-faceted omics analysis

To establish the analytical robustness of AstraKit for somatic mutation profiling, workflows for analyzing variants and mutated genes, mutual exclusivity, and affected pathways enable effective analysis of large patient cohorts (Figure 3A). These analyses identify significantly altered genes, co-occurring mutations, and dysregulated biological pathways. Beyond core genomic analysis, AstraKit integrates multi-omics data to enable functional validation of variant impact and therapeutic target prioritization. For instance, users can stratify cohorts by mutation status (e.g., *KRAS* amplification) to compare transcriptomic or proteomic profiles between mutant and wildtype groups. Critically, AstraKit’s survival modeling workflow executes Lasso-penalized Cox regression using user-specified inputs (e.g., 48 MSigDB Hallmark pathway scores and binary mutation status of the top 50 altered genes), generating risk scores that automatically stratify cohorts for Kaplan-Meier survival analysis (Figure 3B). This enables identification of mutation-pathway signatures prognostic of clinical outcomes while controlling overfitting through crossvalidated regularization.

**Figure 3:**
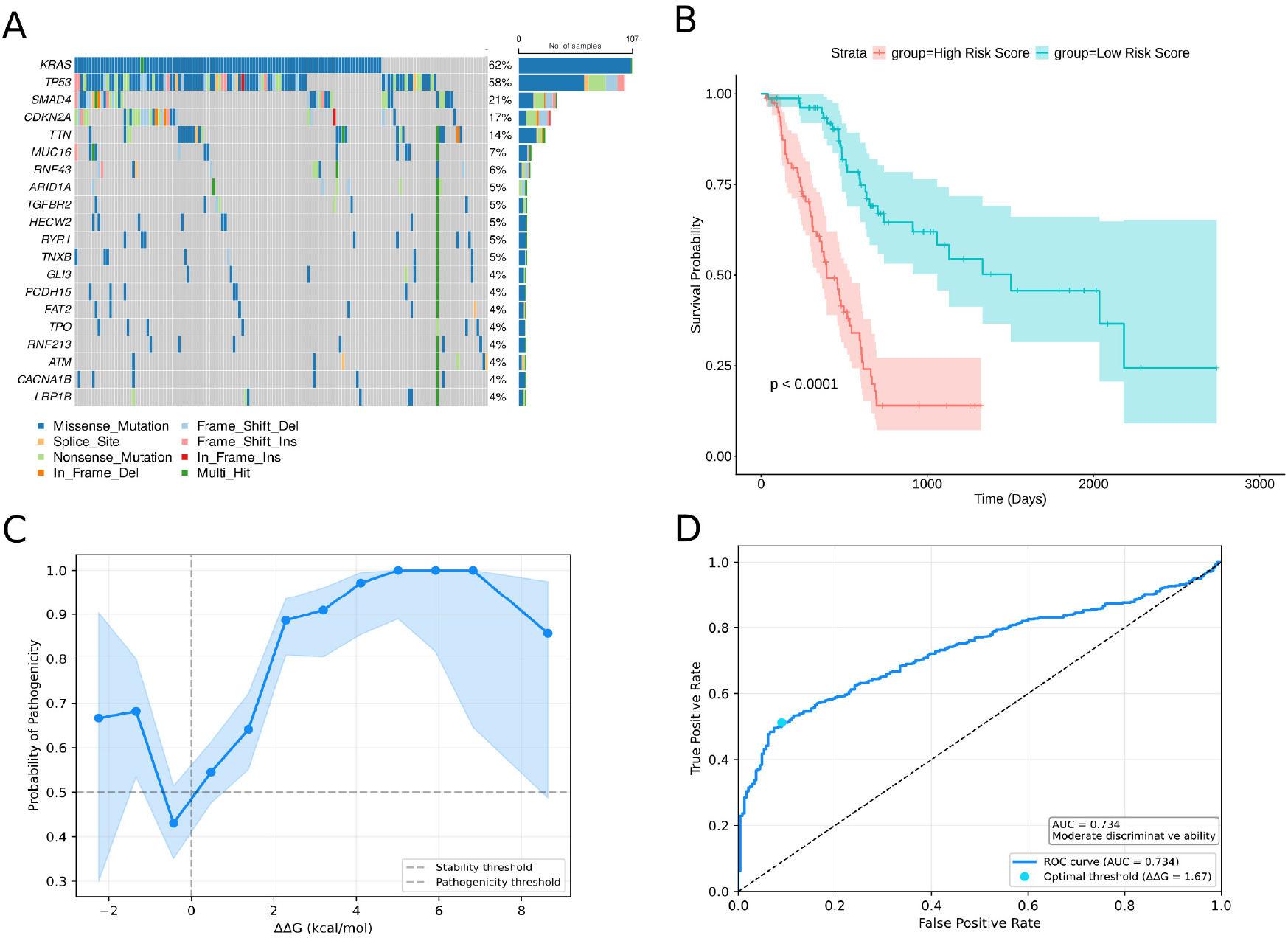
AstraKit variant cohort analysis overview. (**A**) Automated oncoprint analysis of the 20 most recurrently altered genes in the TCGA PAAD (pancreatic adenocarcinoma) cohort, generated directly using AstraKit’s variant cohort analysis workflows. (**B**) Kaplan-Meier curve for overall survival in TCGA-PAAD stratified by risk score from Lasso-regularized Cox model (48 MSigDB Hallmark pathways, top 50 mutated genes with median split). (**C**) Correlation analysis between annotated variant features, including computational predictions of protein stability changes (e.g., using FoldX) and AlphaMissense-predicted pathogenicity scores for somatic TP53 variants. (**D**) Receiver operating characteristic (ROC) between pathogenicity prediction and protein stability change, indicating a modest correlation (AUC = 0.73).

Critically, AstraKit links variant annotation with downstream feature analysis. For instance, protein stability changes (predicted by FoldX) can be systematically correlated with pathogenicity scores, like AlphaMissense. When analyzing TP53 variants, we observed elevated pathogenicity probabilities for both stabilizing (negative ΔΔ*G*) and destabilizing (positive ΔΔ*G*) mutations, suggesting stability-independent pathogenic mechanisms (Figure 3C). A receiver operating characteristic (ROC) analysis using ΔΔ*G* to predict pathogenicity yielded an area under the curve (AUC) of 0.73, indicating a fair correlation between protein stability perturbation and pathogenicity (Figure 3D).

### 3.4 Drug screen validation with clinical outcome

Complementary to molecular profile analysis, AstraKit provides streamlined workflows to analyze targeted therapies for patient cohorts, identifying the most sensitive/resistant drug classes (Figure 4A), drugs (Figure 4B), biomarkers (Figure 4C), and tumor types (Figure 4D). A key innovation is the correlation of preclinical drug screen results with clinical evidence using matched molecular profiles. The associated workflow loads genetic profiles (VCF) and drug screen data (e.g., IC_50_ or GR*max*.), with useradjustable parameters for normalization, cancer type filtering, and clinical evidence sources. To validate this approach, we analyzed GDSC drug screens alongside the matching DepMap mutational profiles of the employed cell lines. We generated cell-line-wise VCF files and mapped their mutational profiles to therapeutic options using Onkopus using exact match searches. This integration reveals drugs with high experimental efficacy (e.g., elevated pIC_50_) but limited clinical exploration (fewer clinical records) (Figure 4E). Conversely, comparing pIC_50_ values with clinical outcomes per cell line highlights models with strong translational potential (Figure 4F). Interactive sunburst visualizations (Figure 5A) enable multiscale exploration of therapeutic options by drug class, biomarker, and tumor type. For TCGA PAAD patients, ROC analysis of cell line-derived predictions versus clinical outcomes showed strong concordance for specific cell lines (COSMIC ID 753595: AUC 0.89; 907285: AUC 0.88) but significant variability across other lines, underscoring the need for context-specific model selection (Figure 5B).

**Figure 4:**
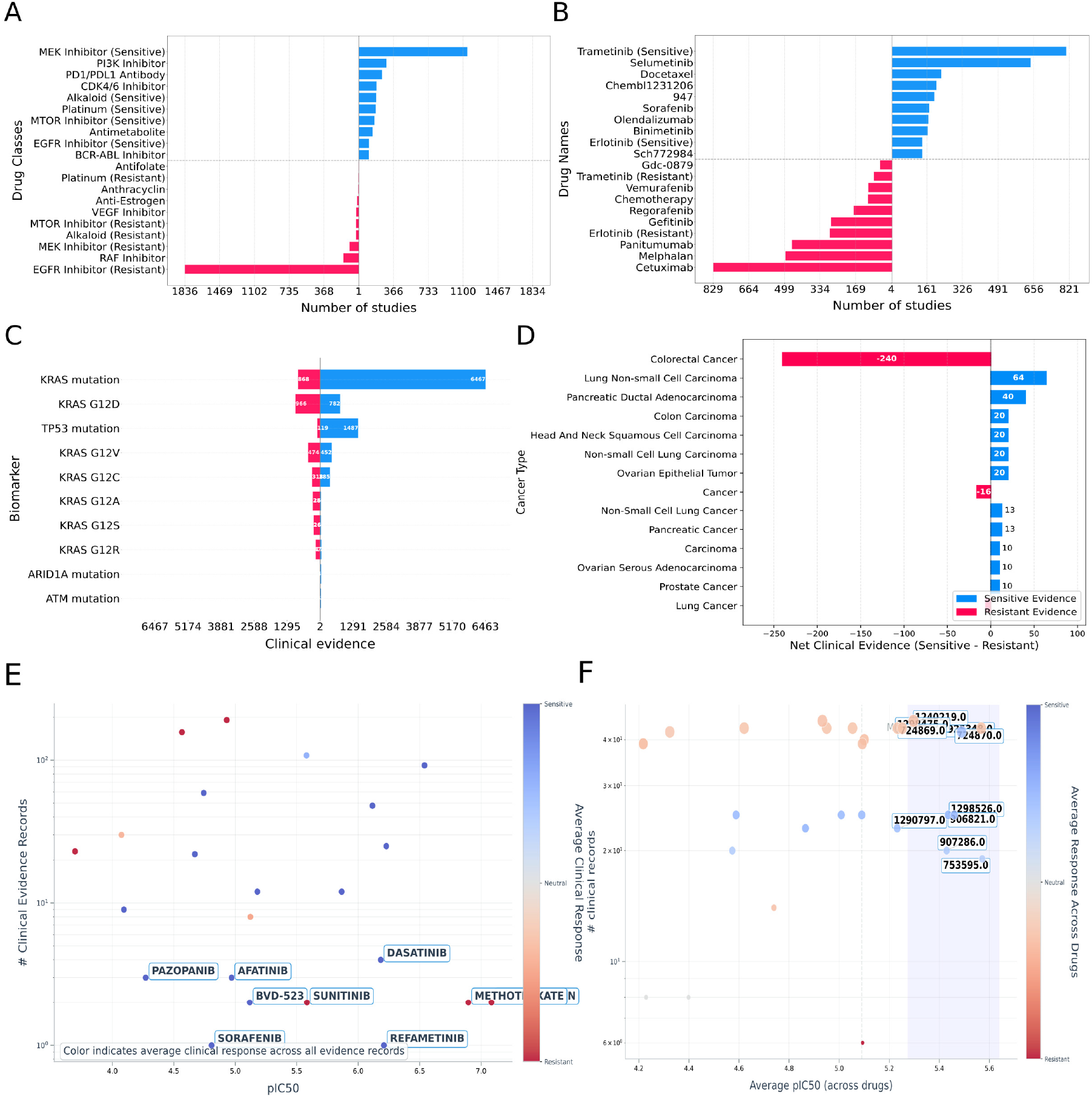
Clinical evidence analysis for patient cohorts and drug screen analysis. (**A**) Top 10 sensitive and top 10 resistant drug classifications for pancreatic cancer patients. Clinical studies were retrieved using Onkopus exact matches, which retrieves only evidence for the exact biomarker found in patient’s mutational profile. (**B**) Top 10 sensitive and resistant therapeutics, and (**C**) number of studies per biomarker with sensitive (blue) and resistant (red) outcome. (**D**) Top sensitive and resistant clinical outcome per cancer type, based on PAAD patient biomarkers. (**E**) Comparison of clinical evidence records and drug screen results (pIC50) per therapeutic option. Drugs with low clinical evidence, but high pIC50 values are highlighted, indicating potentially underexplored, sensitive therapeutics.(**F**) Comparison of clinical evidence and drug screen results based on cell lines.

**Figure 5:**
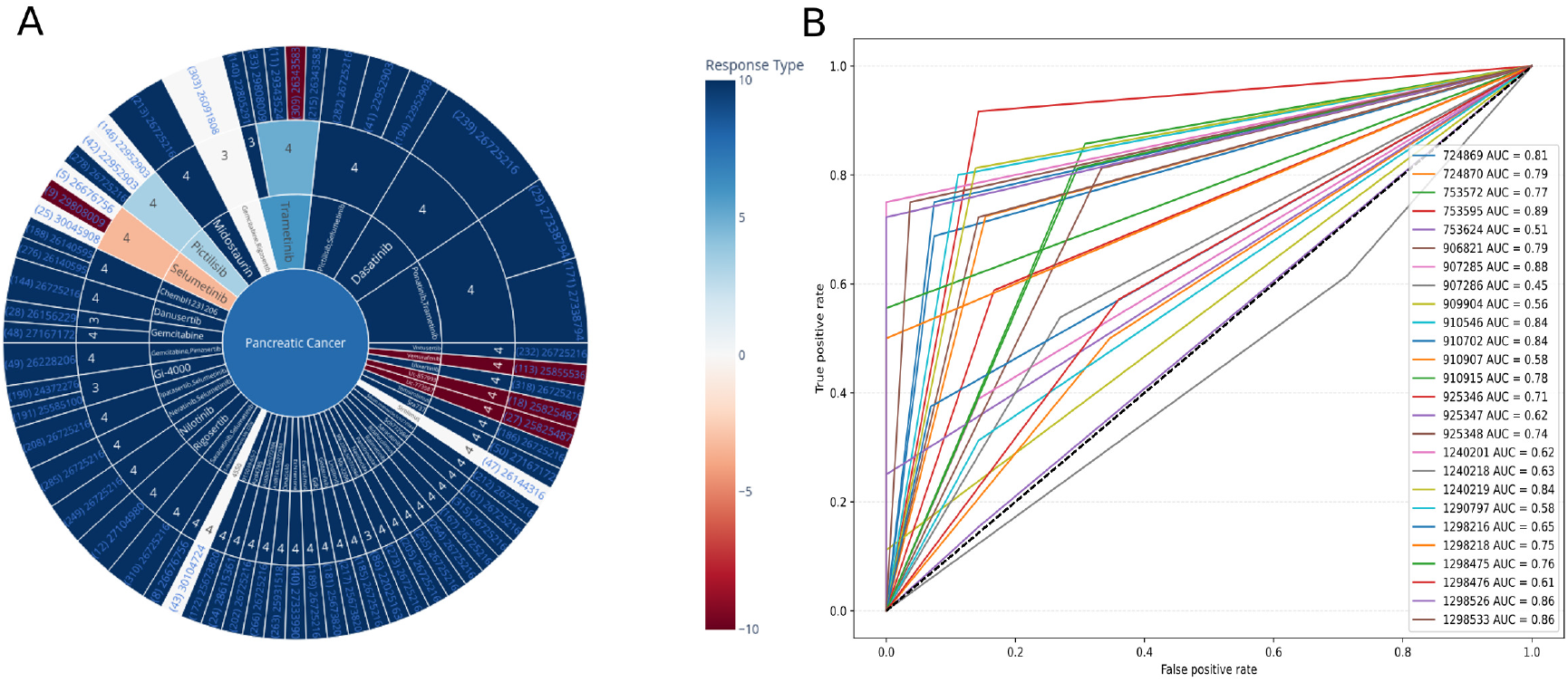
Drug screen analysis: Comparison between experimental drug sensitivity and clinical outcome. (**A**) Clinical studies for KRAS:p.G12D, filtered to clinical evidence for pancreatic cancer, visualized in the Onkopus sunburst visualization. Blue results indicate sensitive, red resistant outcome. (**B**) Receiver operating characteristic of clinical outcome of TCGA PAAD patient biomarkers predicting experimental drug sensitivity (IC_50_) per cell line. The correspondence between drug screen and clinical studies’ results varies significantly across cell lines, ranging from an area under the ROC curve (AUC) from 0.45 up to 0.89.

## 4 Discussion

We presented AstraKit, a comprehensive suite of modular, KNIME-based workflows designed for diverse applications in bioinformatics and precision medicine, including clinical-grade genetic variant interpretation, multi-layered omics analyses, and translational drug-response modeling. Unlike monolithic pipeline frameworks, AstraKit emphasizes visual configurability and end-to-end automation while maintaining compatibility with clinical data governance requirements. As AstraKit leverages the KNIME analytics platform, all workflows operate through graphical, drag-and-drop nodes that eliminate the need for coding expertise. While command-line tools like Snakemake and Nextflow excel in scalable highperformance computing environments, and Galaxy offers cloud-based accessibility, KNIME provides a distinctive balance of visual workflow design, local execution capability, and integration with statistical/ML libraries, making it particularly suitable for processing sensitive clinical data under regulatory frameworks. AstraKit addresses critical gaps in existing tools by replacing error-prone manual tool chaining with fully automated genomic processing pipelines, encompassing variant interpretation, filtering, and coordinate conversion (LiftOver). They further enable scalable cohort analysis for mutational signature profiling, mutual exclusivity testing, and stratified omics investigations. Finally, they integrate clinical evidence with preclinical drug-response data through systematic correlation of patient genomic biomarkers with in vitro drug sensitivity metrics, thereby identifying candidate therapies even when clinical evidence is limited.

A key innovation is AstraKit’s modular architecture for variant processing. Prior frameworks primarily automate variant calling, but offer limited flexibility in downstream annotation and filtering. In contrast, AstraKit decomposes the entire variant interpretation workflow into discrete, interchangeable nodes, allowing users to dynamically configure annotation sources (e.g., ClinVar, GTEx, CIViC), filtering criteria, and LiftOver steps without scripting. For cohort-scale analyses, AstraKit identifies mutational patterns, pathway enrichments, and mutual exclusivity across patient groups. Critically, while most dependencies are auto-installed via KNIME, license-restricted tools (e.g., FoldX for protein stability prediction) require pre-installation by the user, a limitation noted in our documentation.

In drug-response modeling, AstraKit bridges two complementary approaches: First, clinical evidencebased interpretation is enabled by annotating variants to match therapies with existing guidelines (e.g., OncoKB, CIViC). Second, AstraKit provides a preclinical data-driven discovery by correlating genomic alterations with high-throughput drug screen results to uncover novel biomarker-drug associations. This requires rigorous normalization of dose-response curves and statistically robust thresholding (e.g., using FDR correction) to discretize continuous drug sensitivity metrics values into clinically meaningful “sensitive” vs. “resistant” categories, a non-trivial challenge due to cell-line-specific confounders.

Validation of this approach revealed significant variability in drug response correlations across cell lines, suggesting that lineage-specific biomarkers (e.g., tissue-of-origin effects) critically influence sensitivity. This aligns with recent work by Schmidt et al. (2025) [42], who applied this methodology using MTB-Report for retrieving clinical evidence. The comparison to experimental drug sensitivity revealed T-cell malignancies and identified DDR pathway alterations as biomarkers for WEE1 inhibitor sensitivity, demonstrating how preclinical-clinical integration can reveal actionable targets absent from current databases. In summary, AstraKit delivers a unified, clinically adaptable platform covering the full precision oncology workflow, from raw sequencing data to therapeutic recommendations, while ensuring local execution for data privacy and regulatory compliance. By unifying fragmented tools into auditable, visual workflows, AstraKit accelerates biomarker validation and the translation of genomic insights into personalized treatment strategies, particularly benefiting clinicians and experimental biologists.

## Data availability

The KNIME workflows are publicly available at KNIME HUB at https://hub.knime.com/bioinf_goe/spaces/Public/~Zd6-FraRjnV7MMpu/. The source code of the KNIME workflows and the data preprocessing scripts are available at https://gitlab.gwdg.de/MedBioinf/mtb/astrakit. The source code of all Onkopus modules is available at https://gitlab.gwdg.de/MedBioinf/mtb/onkopus, including instructions on how to install the Onkopus modules locally. All modules have been generated from freely available data sources.

## Author contributions

J.D. and N.S.K. developed the conceptualization and methodology. N.S.K., K.K. and M.S. implemented the KNIME workflows. N.S.K. wrote the manuscript. All authors approved the manuscript.

## Acknowledgments

The authors thank the Molecular Tumor Board of the University Medical Center Göttingen (UMG), namely Alexander König, Li Beißbarth and Kirsten Reuter-Jessen, for providing constant feedback and discussions on clinical needs and use cases. The authors also thank the International Max Planck Research School for Genome Science (IMPRS-GS).

